# [^3^H]T-401 binding to monoacylglycerol lipase (MAGL) in the brains of patients and mice with temporal lobe epilepsy

**DOI:** 10.1101/2025.06.24.661388

**Authors:** Sanjay S. Aripaka, Burcu A. Pazarlar, Lars H. Pinborg, Karen Bonde-Larsen, Francois Gastambide, Benny Bang-Andersen, Jesper F. Bastlund, Jens D. Mikkelsen

**Affiliations:** Neurobiology Research Unit, University Hospital Rigshospitalet, Copenhagen, Denmark; Epilepsy Clinic, Neurology Clinic, University Hospital Rigshospitalet, Copenhagen, Denmark; Department of Pathology, University Hospital Rigshospitalet, Copenhagen, Denmark; H. Lundbeck A/S, Valby, Denmark; Institute of Neuroscience, University of Copenhagen, Denmark

## Abstract

Monoacylglycerol lipase (MAGL) inhibitors are considered as drug candidates for epilepsy. In order to determine the level of MAGL and evaluate changes in the epileptic brain, we have validated and used autoradiography and the MAGL radiotracer [^3^H]T-401 on resected temporal neocortex specimens obtained from patients with temporal lobe epilepsy and in brains from mice with chronic reoccurring seizures. Saturation experiments revealed a K_D_ around 4 nM for the human temporal cortex and 7 nM for the mouse brain. In the human brain, binding of [^3^H]T-401 was detected mostly in the grey matter, and in the subcortical white matter in lower amounts. The levels were strongly correlated in the two cortical compartments. The level of [^3^H]T-401 binding in the human temporal cortex varied about a 4-fold among the patients, but was not correlated to either epilepsy duration or the age of the patients. In the epileptic mouse brain, a significant reduction was observed bilaterally in the hippocampus, as well as in other cortical regions, including the temporal cortex. Interestingly, a highly significant negative correlation was seen between MAGL and binding to the translocator protein 18 kDa (TSPO) expressed in glia.

These data support the presence of MAGL in neuronal and non-neuronal cells, and indicate that MAGL levels in the brains of either patients with epilepsy or mice after intra-hippocampal kainite injection are reduced not only in the epileptic zone in the hippocampus, but more widespread in the brain.

## INTRODUCTION

Monoacylglycerol lipase (MAGL) is a cytosolic serine hydrolase that cleaves monoacylglycerols into fatty acids and is responsible for most of the endocannabinoid 2-arachidonoylglycerol (2-AG) degradation (Bertrand et al., 2010; Blankman and Cravatt, 2013; Chanda et al., 2010; Savinainen et al., 2012; Schlosburg et al., 2010). 2-AG serves as an endogenous ligand with agonist effect on the cannabinoid (CB)1 and CB2 receptors (CB2R) and functions as a retrograde neurotransmitter and inhibiting presynaptic neurotransmitter release (Alger, 2002; Bisogno et al., 1999).

An important metabolite also resulting from the degradation of 2-AG by MAGL is arachidonic acid (AA) which then leads to downstream production of proinflammatory eicosanoids (Nomura et al., 2011; Schlosburg *et al*., 2010).

Attempts have been made to develop MAGL inhibitors (Gil-Ordonez et al., 2018) that regulate levels of 2-AG and AA for the treatment of many CNS disorders characterized by excessive neurotransmission and neuroinflammation. Recent results show that such inhibitors have effects not only in animal models of convulsions (Griebel et al., 2015; Sugaya et al., 2016; Terrone et al., 2018; von Ruden et al., 2015; Zareie et al., 2018), but also a number of other CNS disorders (Blankman and Cravatt, 2013; Chen, 2023; Fowler, 2012; Grabner et al., 2017; Mulvihill and Nomura, 2013).

With regard to epilepsy, endocannabinoids directly target hippocampal glutamatergic neurons to protect against acute epileptiform seizures in mice (Monory et al., 2006; Sugaya *et al*., 2016), and 2-AG and subsequent stimulation of CB receptors is considered to underlie the seizure reducing effect of MAGL inhibitors (Rocha et al., 2020).

The cellular substrates mediating the action of MAGL inhibitors are not fully understood, and as to whether MAGL expression and activity are changed in chronic epilepsy have not been investigated. In the present study, we aimed to measure the level of MAGL in the brains of patients with temporal lobe epilepsy and mice injected with kainic acid in the hippocampus as a model of this disease using a novel radiotracer T-401This radioligand can be displaced by pretreatment with the MAGL inhibitor JW642, and is considered suitable for evaluations of target engagement in the primate CNS (Hattori *et al*., 2022) and has together with a derivative been used in positron emission tomography (PET) in rat, monkey and human (Hattori *et al*., 2019; Hattori *et al*., 2022; He *et al*., 2023; Takahata *et al*., 2022). In line with these observations, we tritiated T-401 to determine binding levels as a marker of MAGL concentrations in the brain. This investigation was focused on eventual changes in binding levels in various regions of the epileptic brains either obtained from neurosurgical resections from patients with temporal lobe epilepsy or from mice after status epilepticus and epileptogenesis.

## MATERIALS AND METHODS

### Radioligands and compounds

T-401 ((*R*)-1-(3-(2-fluoro-4-methylpyridin-3-yl)phenyl)-4-(4-(thiazole-2-carbonyl) piperazin-1-yl)pyrrolidine-2-one) (Hattori *et al*., 2019) and ABD-1970 (1,1,1,3,3,3-hexafluoropropan-2-yl-4-(2-(8-oxa-3-azabicyclo[3.2.1]octan-3-yl)-4-chlorobenzyl) piperazine-1-carboxylate) (Clapper et al., 2018) were synthesized by H. Lundbeck A/S. T-401 was used for tritium labeling using Crabtree’s catalyst for the direct H/T-exchange; Specific activity: 52.8 Ci/mmol (1954 GBq/mmol) obtained from Tritec AG, Teufen, Switzerland. Also, [^3^H]PBR-28 was synthesized and obtained from Tritec AG, Teufen, Switzerland; Specific activity: 80.7 Ci/mmol (3064 GBq/mmol).

### Patients included in the study and tissue processing of resections

Human neocortex samples were obtained from drug-resistant temporal lobe epilepsy patients undergoing temporal lobe resection surgery at the Department of Neurology and Neurosurgery at Rigshospitalet, Copenhagen. The study was approved by the Ethical Committee in the Capital Region of Denmark (H-2-2011-104) and written informed consent was obtained from all patients before surgery. All patients included exhibited seizure semiology, imaging, and ictal EEG findings concordant with an epileptogenic zone in the medial temporal lobe seizures. The temporal neocortex tissue is removed to reach the epileptogenic zone. A total of 19 samples were included in the study (10 females and 9 males, age between 18 and 64 years; average 39.5 years, duration of disease was between 4 and 47 years).

After resection of the temporal cortex, the tissue was briefly washed in physiological saline transported to the laboratory and frozen in crushed dry ice and stored at -80 °C until further processing. Frozen specimens were placed in a cryostat and 12 µm serial sections that included both grey and white matter were collected. The frozen sections were placed in duplicates on Superfrost precoated slides (Thermo Scientific).

### Animals and tissue processing

A total of 14 C57BL/6J male mice (Janvier, France) aged between 10 and 12 weeks were injected with kainic acid (KA) (50 nL of a 20 mM KA solution dissolved in 0.9% NaCl solution) (Sigma-Aldrich, Lyon, FR,) into the right dorsal hippocampus using stereotaxic techniques (coordinates: AP = −2.0 mm, ML = −1.5 mm, DV = −2.0 mm from bregma) under general anesthesia with isoflurane. Similarly, a batch of age matched C57BL/6J male mice (Janvier, France) were stereotaxically injected with saline (n = 8) and used as SHAM group and another batch of C57BL/6J male mice at the same age (n = 8) were used as naïve group.

For electroencephalography (EEG) the KA and SHAM groups were under the same surgery and anesthesia implanted with a bipolar electrode, made of two twisted polyester-insulated stainless-steel wires, into the right dorsal hippocampus using the same stereotaxic coordinates as for KA injection with reference electrodes in left and right frontoparietal cortex and the cerebellum. The five electrodes were then soldered to a female connector and fixed to the skull with dental cement. After this procedure, the animals were kept in individual cages and underwent an approximate 3-week epileptogenic period before reaching a stable level of spontaneous recurrent hippocampal paroxysmal discharges (HPD).

At the end of this phase, a control EEG recording was performed using System Plus Evolution (Micromed France, Mâcon, France, 512 Hz sampling rate, low pass filter: 150 Hz, high pass filter: 0.008 Hz). The overall quality of the EEG signal and presence of HPDs were evaluated, and a cohort of 13 KA injected C57BL/6J mice were selected as fulfilling the minimal criteria of recurrence ˃20 HPD/hour with minimal duration of HPD ˃ 5 sec (thus, one animal was excluded due to lack of EEG signal).

All the animals in this study were killed at the age of 44 weeks and their brains were removed and frozen in crushed dry ice and stored at -80ᵒC until they were sectioned.

From the frozen brains, 12 μm coronal sections were cut on a cryostat (Cryostar NX70, Thermo Scientific) and placed on SuperFrost pre-coated slides (Thermo Scientific). The anatomical regions of interest were the cingulate cortex, (Bregma (B) 1.42 mm, Interaural (I) 5.22mm), striatum (B 0.02 mm, I 3.82 mm), dorsal hippocampus (B -1.94 mm, I 1.86 mm) and ventral hippocampus (B -3.16 mm, I 0.64 mm) and one section from each region was mounted on the glass slide.

### Autoradiography for [^3^H]T-401 and [^3^H]PBR28

The glass slides with either human or mouse brain sections were collected separately. These set of slides were exposed in the same autoradiography experiment to reduce inter-assay variation. First, glasses were allowed to reach room temperature, and then pre-incubated in buffer (50 mM Tris-base, 1.5 mM MgCl_2_, 0.5% bovine serum albumin (BSA), (pH 7.4) twice for 10 minutes. Slides were then incubated for 120 min under constant shaking in buffer (50 mM Tris-base, 140 mM NaCl, 1.5 mM MgCl_2_, 5 mM KCl, 1.5 mM CaCl_2_ and 0.5% BSA) with [^3^H]T-401. For the saturation study, duplicate sections from either two different patients or from one experimental mouse were incubated in concentrations spanning from 3.1nM to 100nM of the radioligand, and non-specific binding was determined in the presence of 10 μM of cold ligand itself or 10 μM ABD-1970. Based on the saturation experiments in mouse and human, we used a fixed concentration at 7.5 nM [^3^H]T-401 for the human, and 25 nM [^3^H]T-401 for the mouse for displacement and comparative studies.

For all autoradiography experiments with [^3^H]T-401, slides were washed for 3 × 5 min with ice cold pre-incubation buffer followed by a quick dip in ice cold distilled water, air-dried and placed overnight in a paraformaldehyde vapor chamber at 4 ℃. After fixation, the glass slides were kept in a silica gel desiccator for 45 min to remove any leftover moisture before exposing them to the imaging plate. Then, glass slides were put together with tritium standards ([^3^H]microscale ART0123 (0-489.1 nCi/mg) and ART0123B (3-109.4 nCi/mg) American Radiolabeled Chemicals, Inc., St. Louis, USA) in a radiation-shielded imaging plate cassette (Fuji casette2 BAS-TR 2040, Fujifilm, Tokyo, Japan) with a tritium-sensitive imaging plate (Fuji IP BAS-TR 2040, Fujifilm, Tokyo, Japan). The glass slides were exposed to image plates for 2 days at 4℃. After exposure, the imaging plate was scanned using Amersham™ Typhoon™ IP Biomolecular Imager (GE healthcare, Chicago, USA) at a pixel size 25µm.

Mouse brain sections adjacent to those exposed for [^3^H]T-401 were also applied for autoradiography for [^3^H]PBR-28. The sections were preincubated in the same buffer for 10 min, and then 1h under constant shaking in buffer (50 mM Tris−HCl, 0.5% bovine serum albumin (BSA), 5 mM MgCl_2_, 2 mM EGTA) with 3nM [^3^H]PBR-28.

### Quantification of the autoradiograms

As the specimens were coded the investigators were blinded to the nature of the samples, and the images were analysed by two independent investigators. For human sections, the region of interest involved the full extent of the grey and white matters in two separate tissue sections. For the mice sections, the regions of interest were defined based on specific landmarks in the sections. Statistical analysis was performed using GraphPad Prism version 9.1.0. The overall effect on treatment was assessed using a Student t-test analysis. The correlation of binding levels with another variable was assessed using the Pearson correlation coefficient. A *P* value of < 0.05 was considered statistically significant for all comparisons. Data are expressed as mean ± SD.

## RESULTS

### Validation of the radiotracer [^3^H]T-401 in human brain tissue

Serial frozen sections from temporal cortical resections obtained from two patients with temporal lobe epilepsy were used in preliminary experiments to validate the binding characteristics of the MAGL radiotracer [^3^H]T-401. Because autoradiography using [^3^H]T-401 has not been reported before, preliminary validation experiments were carried out to determine the optimal temperatures and times for the incubation and washing steps. Room temperature was found to be optimal for the incubations, and cold (4 °C) was found to be optimal for the washing steps. Different incubation and washing times were also tested. Based on these experiments the autoradiography was carried out with 120 min incubation time at room temperature, and the washing for a total of 15 min at 4 °C. Repeated analysis revealed an inter-assay variation of 3.2 %.

### [^3^H]T-401 binding could be saturated and displaced in the human brain

In the human temporal cortex, the radiotracer bound predominantly to the grey matter, but relatively weak specific binding was also detected in the subcortical white matter (Fig. 1).

**Fig. 1.**
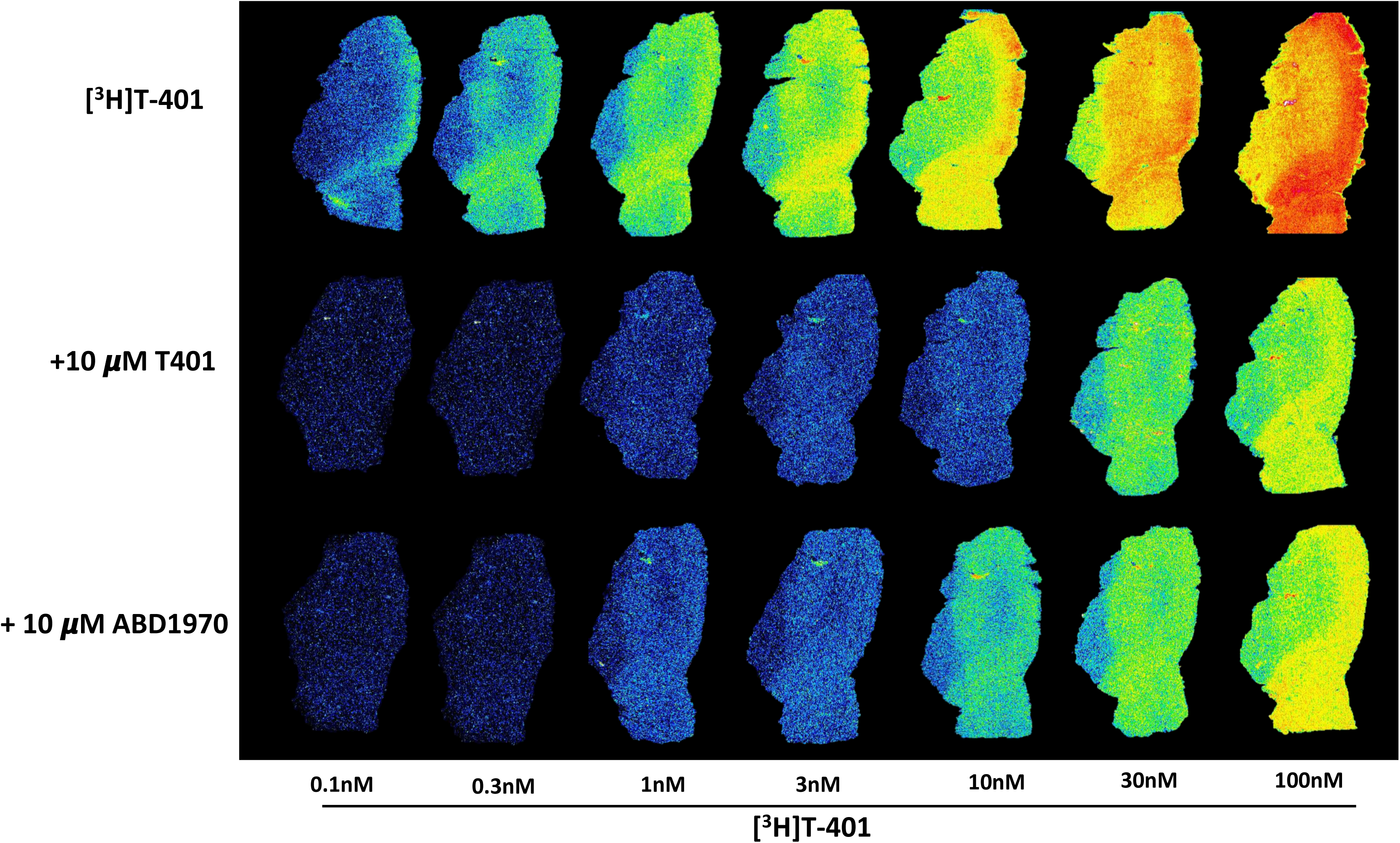
Representative autoradiograms of serial sections obtained from a human temporal cortex resection from one treatment resistant temporal lobe epilepsy patient. The serial sections from this specimen have been incubated together with increasing concentrations of [^3^H]-T401 (A). Displacement with the cold ligand T-401 (10 µM) shows complete block up to 10 nM (B), whereas displacement of the ligand with 10 µM ABD1970 showed only complete block up to 3 nM of the ligand (C) The highest density is observed in the grey matter, with the most in the superficial layers of the cortex. Low labelling is also observed in the white matter.

Image analysis and standard corrected quantification revealed that binding of [^3^H]T-401 reached saturation in the human brain tissues in low nM concentrations (Fig 2A). The saturation experiment with the radioligand [^3^H]T-401 in the presence or absence of high concentrations of the displacement ligands (T-401 and ABD1970) (see Figs. 1 and 2A, 2B) revealed a K_D_ at 4.7 nM ± 0.8 nM as a mean from two independent experiments in two patients. No differences in K_D_ values were seen between regions.

**Fig. 2.**
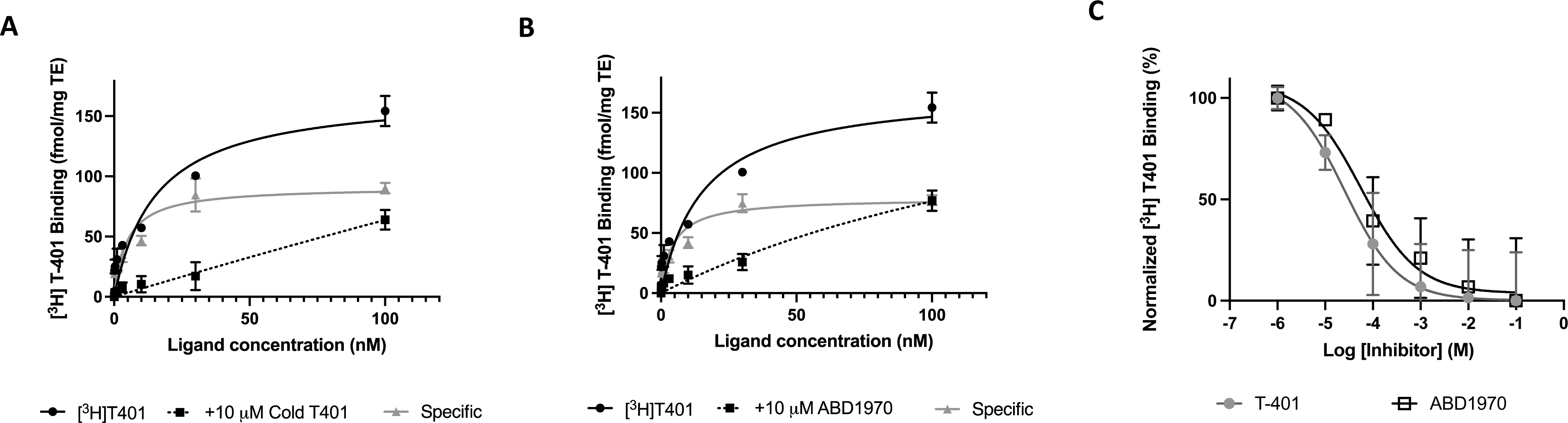
Saturation curves with and without co-incubation with a high concentration (10 µM) of either the cold ligand T-401 (A) or another structurally different MAGL inhibitor ABD1970 (B) show that [^3^H]-T401 is saturable and displaceable in the human cortex. The analysis revealed a saturation and the K_D_ was calculated to be 4.7 (cold ligand) and 4.3 (ABD1970). Displacement with 25 nM [^3^H]-T401 in the presence of increasing concentration of the two cold ligands revealed full saturation, and calculated IC_50_ values, 32 nM for T-401 and 63 nM for ABD1970.

A displacement experiment was carried out using the same two MAGL modulators (Fig. 2C). Exposing human samples with the radiotracer at a concentration with full saturation and co-incubation with increasing concentrations up to 100 µM of either the cold ligand or ABD-1970 revealed a sigmoid displacement curve in both cases and the calculated best fit for IC_50_ was found to be 32 nM for T-401 and 63 nM for ABD-1970 (Fig. 2C). Both ligands were able to reach full displacement of [^3^H]T-401 at concentrations up to 1μM.

### Binding in the patient population: Comparison to age and disease duration

Individual sections taken from resections of the temporal cortex from 19 patients were exposed to autoradiography with 7.5 nM [^3^H]T-401, with and without ABD1970. Total, displaced and specific binding were determined in both grey and white matter (Fig. 3A-C). The specific labeling (B_max_) was measured for each patient sample. In the cohort, the level of specific binding was measured to be 27.9 ± 14 fmol/mg tissue equivalent in the grey matter and 10.2 ± 6.1 fmol/mg tissue equivalent in the white matter.

**Fig. 3.**
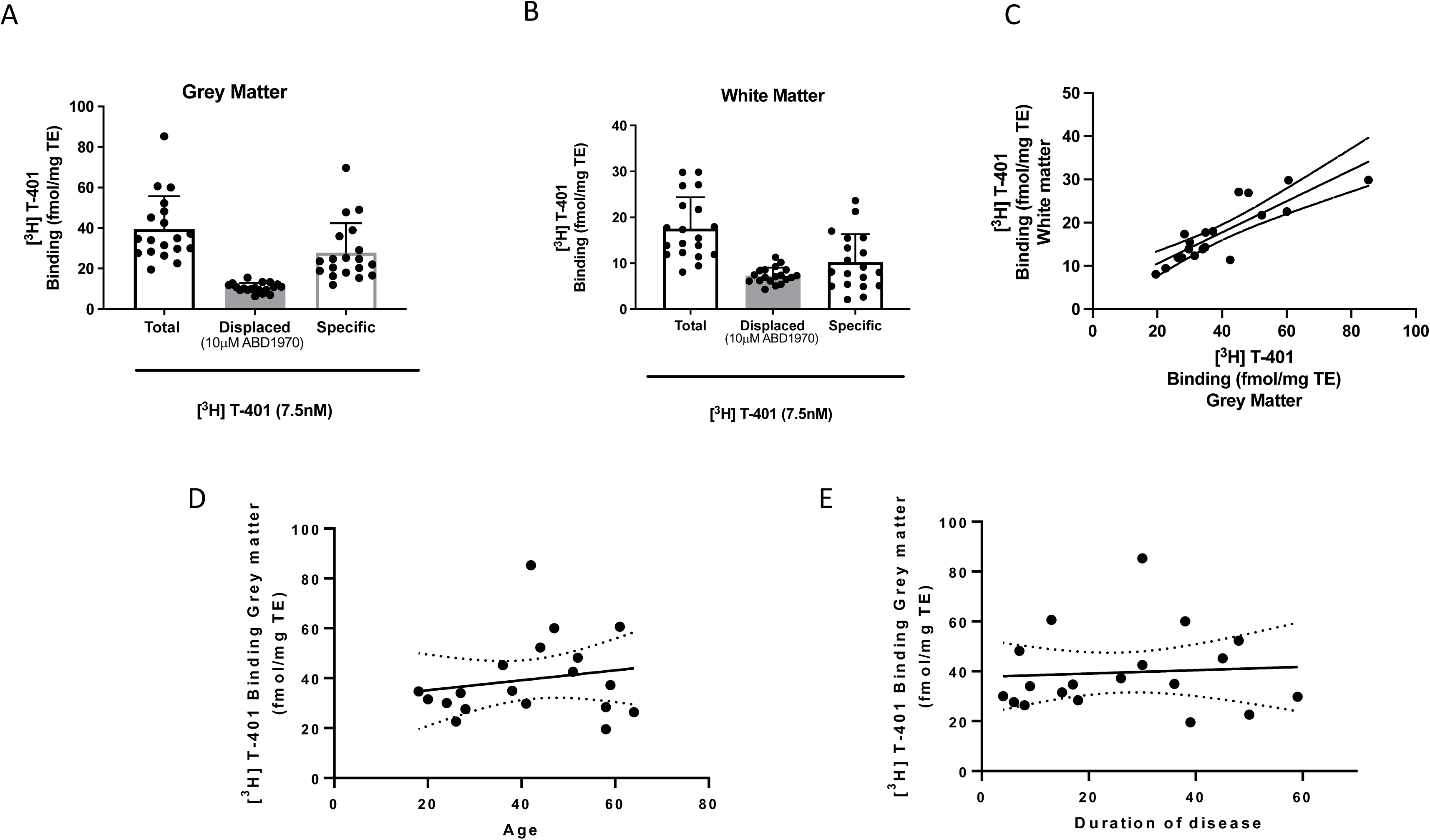
Quantification of total, displaced and specific binding in the temporal cortical resection from patients with temporal lobe epilepsy (n = 19). The total and the specific binding of [^3^H]-T401 in the cohort is found to be rather variable in both the grey (A) and white matter (B). By contrast, little variation is found in the non-specific binding in both regions. Strong correlation (r = 0.85; P < 0.001) is found between the binding in the white and grey matter (C). There is no correlation between the specific binding and age (r = 0.18; p = 0.45) (D), nor duration of disease (r = 0.07; p = 0.77) (E).

Among the patient samples, there was a large variability in the total binding and because the variability in the displaced non-specific binding data was very low the variability is reflected in the specific binding between patients. When correlating the specific binding in the white and grey matter between the patients, a very strong correlation was observed (r = 0.85; P < 0.001) (Fig. 3C), which led us to further analyse any association to clinical and demographic variables. However, no association to the binding levels was found neither to age (r = 0.18; P = 0.45) (Fig. 3D) nor to the disease duration (r = 0.07; p = 0.77) (Fig. 3E) at the time of collection of the samples. Also, the time of collection was not found to affect the binding (not shown).

### [^3^H]T-401 saturation and displacement experiments in the mouse brain

In the mouse brain, the radiotracer [^3^H]T-401 predominantly bound to the grey matter i.e., in the cerebral cortex, hippocampus and thalamus (Fig. 4).

**Fig. 4.**
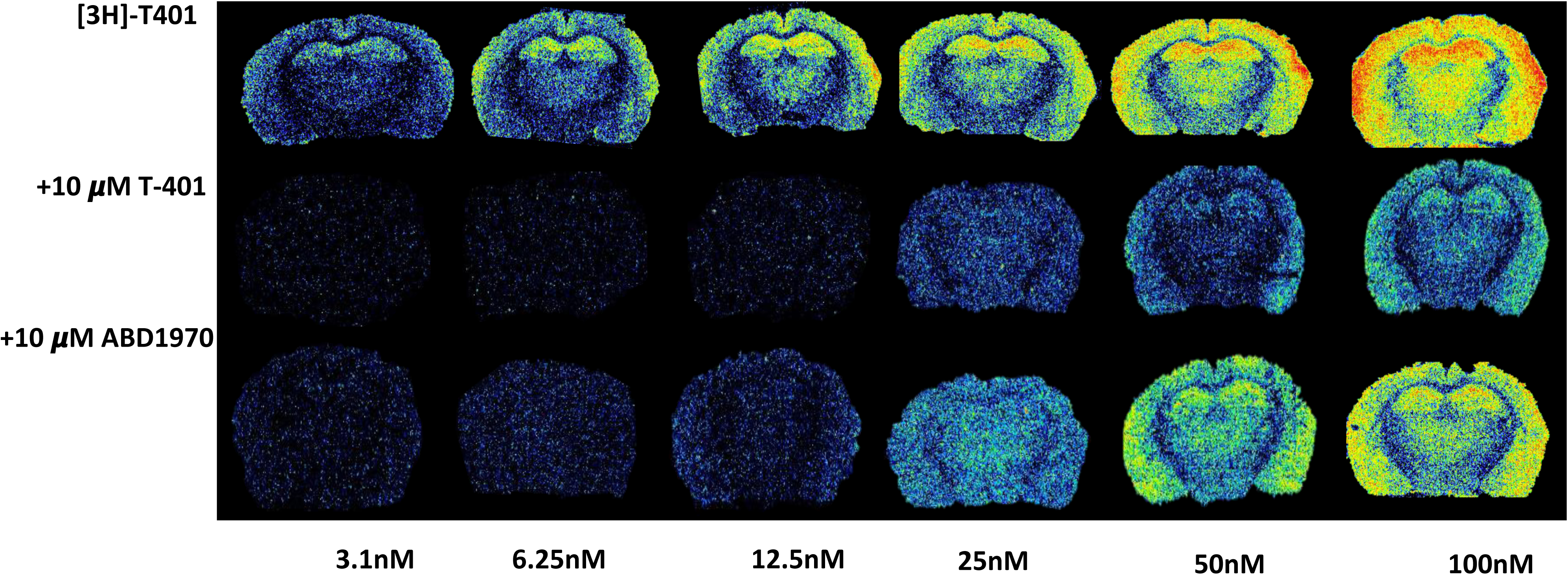
Representative autoradiograms of frontal sections from a mouse brain incubated with increasing concentrations of [^3^H]T-401 (A). The highest density is observed in the grey matter, such as the hippocampus. In the cerebral cortex more labeling is present in the superficial layers and in the parietal and frontal parts, in contrast to the piriform cortex. Displacement with the cold ligand T-401 (10 µM) shows complete block up to 10 nM (B), whereas displacement of the ligand with 10 µM ABD1970 showed only complete block up to 3 nM of the ligand (C).

As illustrated in Fig 4A, incubation with [^3^H]T-401 at increasing concentrations reached saturation in the cerebral cortex and the hippocampus. Co-incubation with 10 µM of either the cold ligand (Fig. 4B), or another MAGL inhibitor ABD-1970 (Fig. 4C) produced in both cases a block in binding up to 25 nM [^3^H]T-401. Specific binding and saturation could also be detected in the hypothalamus and the corpus callosum (white matter), but achieving lower B_max_. Notably, the non-specific binding as revealed with ABD1970 displacement was linear up to 50 nM [^3^H]T-401, but not at 100 nM probably reflecting a low affinity non-specific binding site (Fig 5B, D).

**Fig. 5.**
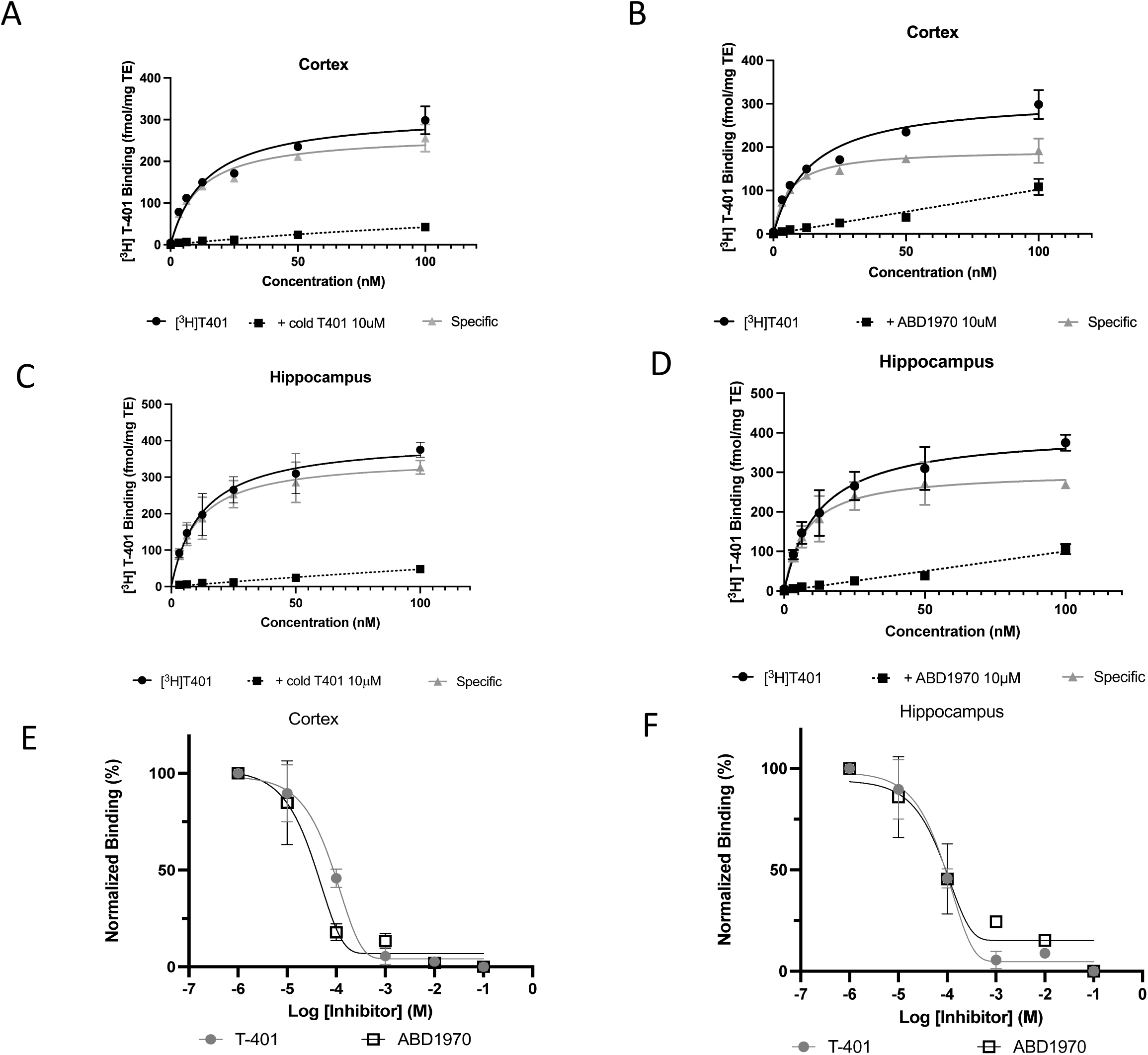
Quantification of binding intensities in different regions of the mouse brain such as the hippocampus (A, C); cerebral cortex (B, D) at increasing concentrations of [^3^H]-T401. The level of specific binding (Δ) is defined by subtracting the non-specific binding (▪) from the total binding (●) after co-incubation with 10 µM T-401 (A, C) or 10 µM ABD1970 (B, D). Inhibition of 25 nM [^3^H]-T401 with increasing concentration of the two cold molecules (T-401 ●; ABD1970 ▪) revealed an inhibition curve in the hippocampus (E) and the cerebral cortex (F). The B_max_ and K_D_ were calculated to be 194 fmol/mg tissue and 5.8 nM for the cortex and 302 fmol/mg tissue and 4.9 for the hippocampus. IC_50_ was calculated to be 41 nM for T-401 and 14 nM for ABD1970.

A displacement study was carried out using the same two MAGL modulators at the concentration of 25 nM [^3^H]T-401 (Fig. 5E, F). Displacement of the radiotracer with increasing concentrations of either the cold ligand or ABD-1970 revealed a calculated best fit and IC_50_ was calculated to be 41 nM for T-401 and 14 nM for ABD1970 as an average of two binding experiments. In contrast to the human brain, full displacement could not be achieved in the mouse brain (Fig. 5E, F).

### Level of [^3^H]T-401 binding in the brain from mTLE mouse model

The binding levels of [^3^H]T-401 in brain sections were analyzed from 3 groups of mice (i.e., naïve, sham, and KA) (Fig. 6). Sections from all animals were incubated together with 25nM of [^3^H]T-401, each slide containing four regions from rostral to caudal (anterior cingulate to ventral hippocampus) (Fig. 6). The KA injection caused a clear and visible loss of [^3^H]T-401 binding in the ipsilateral hippocampal region when compared to the contralateral side (Fig. 6, 7A), but only in the hippocampus. Furthermore, in this brain region not only was there a difference between the two sides, but the level of binding was significantly lower than sham in both the ipsi- and the contralateral side (Fig. 7A).

**Fig. 6.**
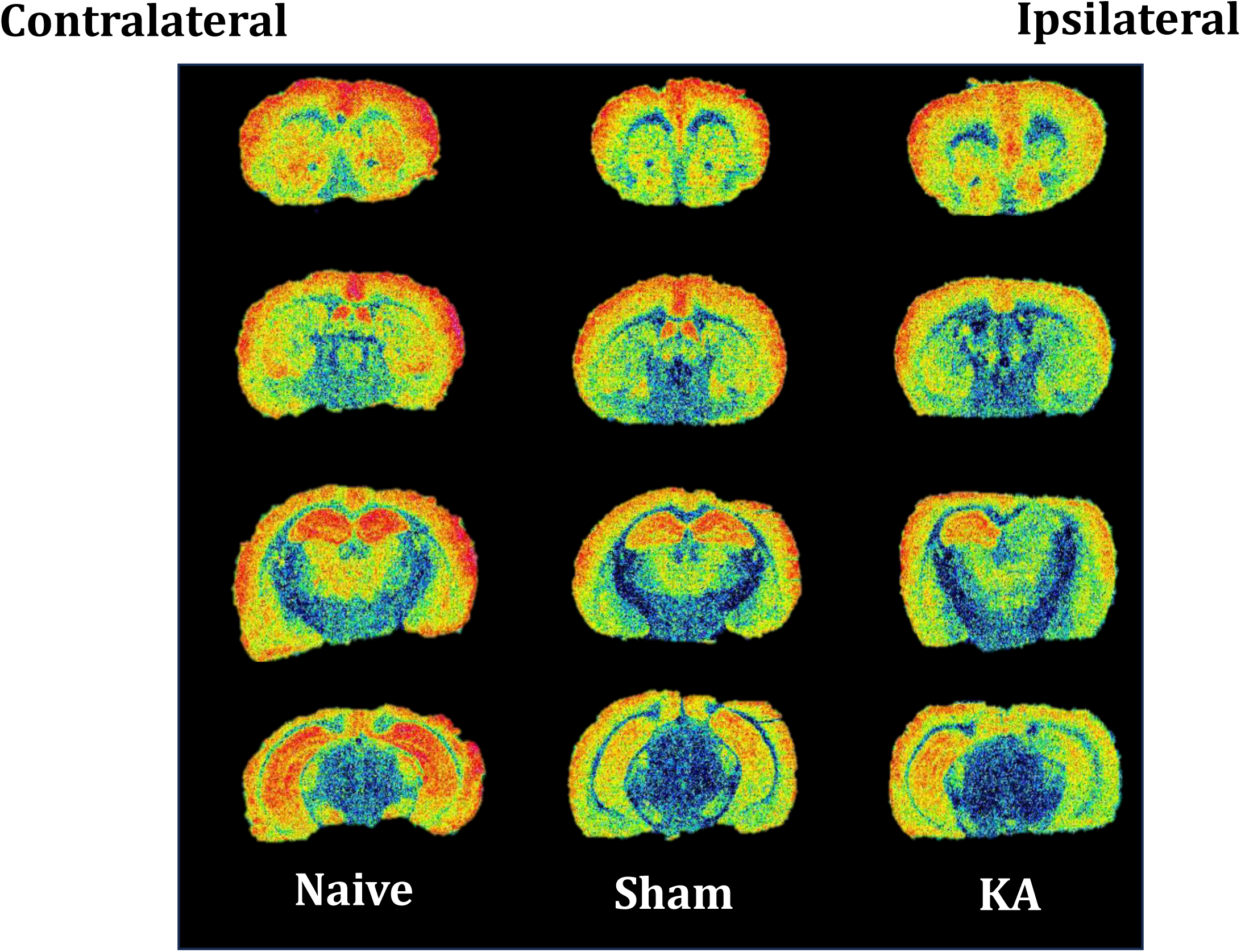
Representative autoradiograms from mice at four different rostro-caudal levels (top to bottom) from the same animal that is either naïve, sham operated, or injected with KA (ipsilateral) All sections were incubated together so the binding density is directly comparable between animals. Note that in the the ipsilateral hippocampus, a lower level of binding can be observed.

**Fig. 7.**
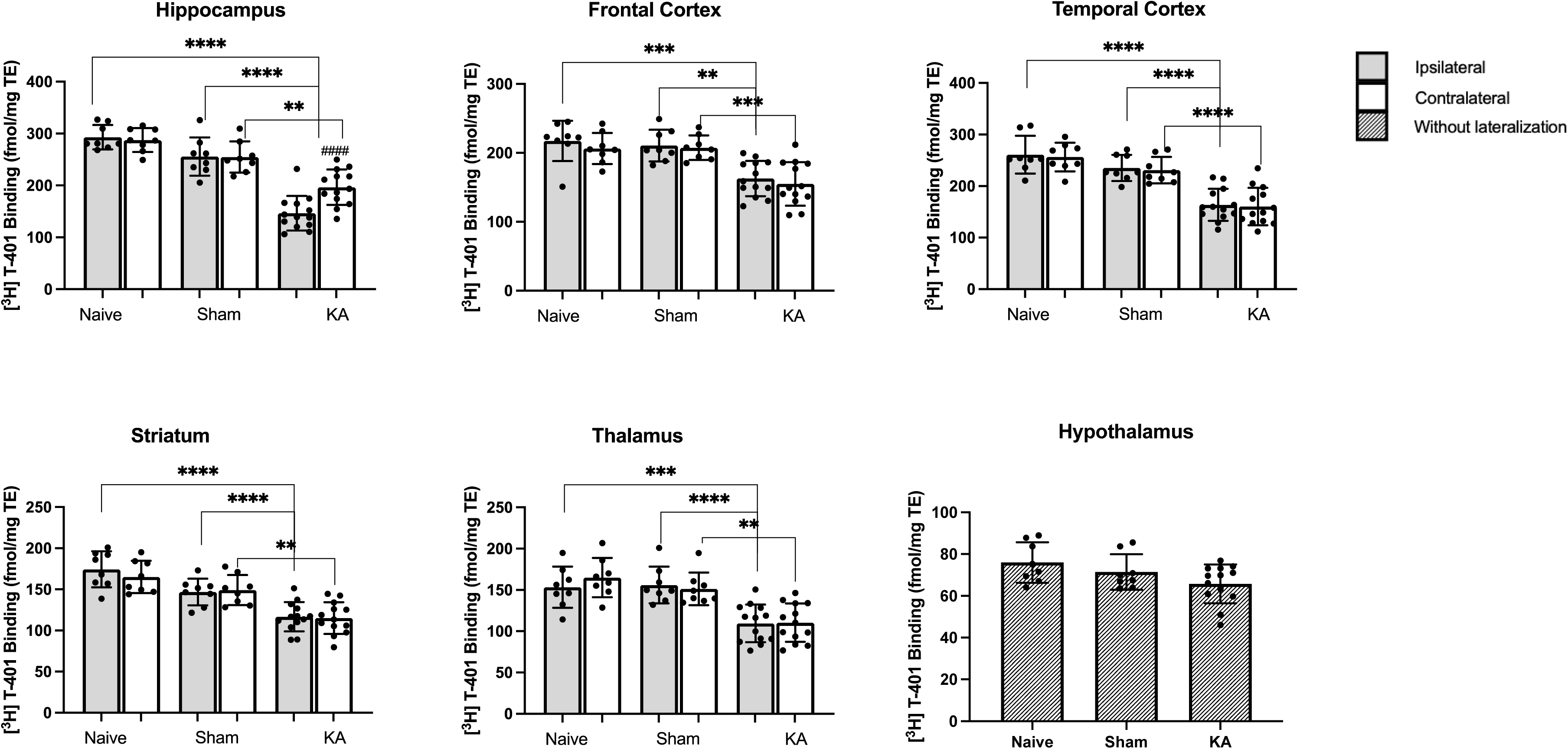
Quantitative estimates of the level of binding in various parts of the brain after chronic seizures and spontaneous excitations. While there was no difference between naïve and sham in any region, a significant reduction was present in the hippocampus (A), frontal cortex (B), temporal cortex (C). striatum (D), and thalamus (E), whereas no difference was observed for the hypothalamus (F). **** P < 0.001 for both sides; #p < 0.01 between the two hemispheres

Next, the [^3^H]T-401 binding in other brain areas were compared to sham control (Fig. 7A-F). Analysis of cerebral cortical subregions represented in the tissue sections revealed a reduction in all parts of the neocortex. For example, a significant decline in [^3^H]T-401 binding in both temporal (Fig. 7B) and in the frontal cortex (Fig. 7C) was significantly lower at both sides.

Subcortical structures, such as the striatum and thalamus also displayed a significant reduction in binding in the KA group (Fig. 7D, 7E). By contrast, binding in the hypothalamus was not affected by the KA injection (Fig. 7F).

The association between [^3^H]T-401 binding and inflammatory status revealed with the in-house validated TSPO tracer [^3^H]PBR28 was also investigated. As expected, a highly significant increase in [^3^H]PBR28 binding was found in the ipsilateral side compared to contralateral side at hippocampal level and the binding level was significantly larger when compared to naïve and sham groups at both sides (Fig. 8A). By contrast, no change was seen in the TSPO binding outside the hippocampus such as the temporal cortex, as well as the frontal cortex (not shown). Interestingly, a significant negative correlation (r = -0.8; p = 0.009) was found between the binding of [^3^H]PBR28 and [^3^H]T-401 in the ipsilateral hippocampus (Fig. 8B), but not in the temporal cortex (Fig. 8C).

**Fig. 8.**
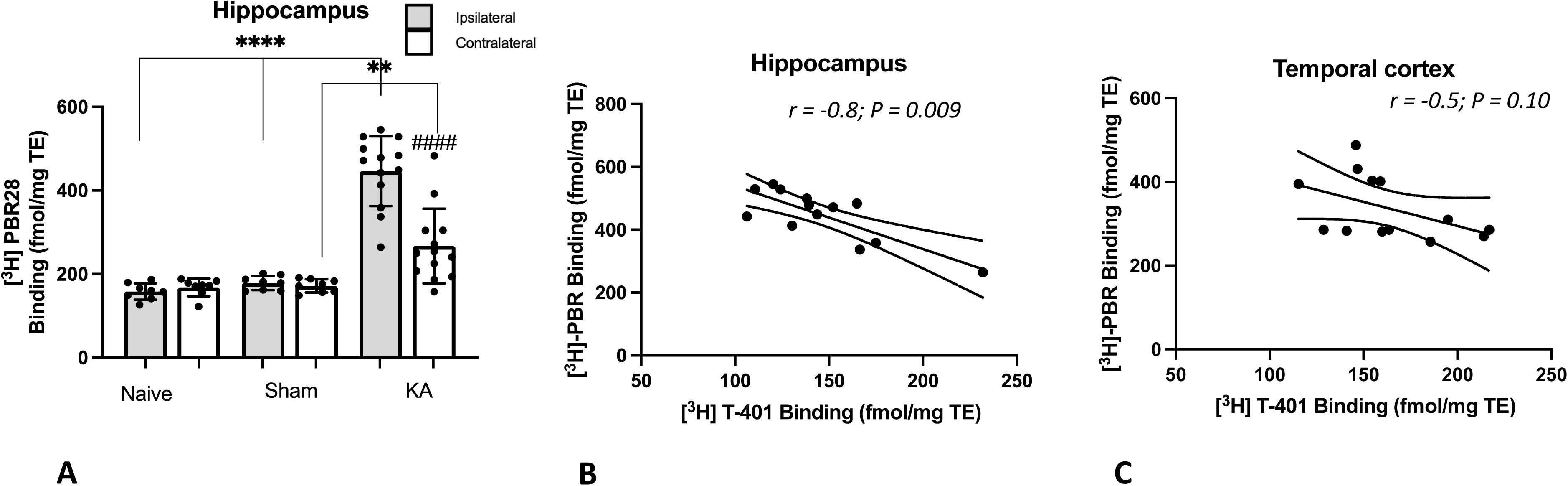
Determination of the level of TSPO binding showed a highly significant increase on both sides (A). Correlation between the level of MAGL and TSPO binding in animals with chronic seizure. Comparison of the binding in TSPO and MAGL revealed a highly significant negative correlation (r = -0.8; P=0.009) in the ipsilateral hippocampus (B), whereas no such correlation was observed for the ipsilateral temporal cortex (r = - 0.47; P = 0.10) (C).

## DISCUSSION

Identification of MAGL mRNA almost completely overlap with the distribution of binding sites presented here (Dinh et al., 2002). In particular, the dominant expression in the cerebral cortex and hippocampus is observed in both *in situ* hybridization and receptor binding.

Binding experiments performed in both human and mouse brain tissues showed that [^3^H]T-401 binding could be saturated at least up to a concentration of 50 nM. These findings supports earlier work that [^3^H]T-401 is a reliable radiotracer for measuring MAGL binding in both primate and rodent brain tissues ( Hattori *et al*., 2019; Mikkelsen et al., 2024). The K_D_ value was low in human brain, higher in mouse brain, and highest in the rat brain ( Mikkelsen *et al*., 2024), perhaps reflecting different concentrations of 2-AG between species (Walter et al., 2004).

Specific binding of [^3^H]T-401 was detected in low levels in the subcortical white matter indicating the presence of MAGL in non-neuronal cells. Expression of MAGL in non-neuronal tissues has been demonstrated by immunolocalization (Uchigashima et al., 2011; Walter *et al*., 2004) and after genetic ablation in glia tissues (Chen, 2023; Zhu et al., 2023). MAGL expression outside neurons occurs in astrocytes (Viader et al., 2015; Viader et al., 2016), and perhaps oligodendrocytes (Gomez et al., 2010). Considering that neurons and astrocytes are the main cellular source of MAGL, it is interesting to speculate whether the changes in MAGL binding occur in one cell type or the other, or both. The relative contributions of neurons and glia to the termination of 2-AG dependent functions, as well as the potential for crosstalk between these cell types in homeostasis of endocannabinoid pools remain largely unknown. The present analysis of MAGL levels in temporal cortex in a cohort of epilepsy patients demonstrated a variability in the binding of [^3^H]T-401 in both the grey and white matter suggesting a heterogeneity of MAGL expression among patients with temporal lobe epilepsy rather than between structures in the same patient. Astrocytes are recognized as integral cellular components and regulators of neurotransmission (Halassa and Haydon, 2010). Thus, 2-AG release may affect CB2 receptors and thereby produce anti-inflammatory effects as seen in a number of models (Du et al., 2011; Panikashvili et al., 2006; Panikashvili et al., 2001; Zhang and Chen, 2008). When eliminating MAGL in only astrocytes LPS-induced effects on prostaglandin E_2_ and D_2_ production is reduced (Viader *et al*., 2015). Further, inactivation of MAGL exclusively in astrocytes exerts upregulated expression of immunogens in microglia (Zhu *et al*., 2023). Microglia which are resident immune cells of the CNS, also contribute to neurotransmission through activity-dependent remodeling and pruning of synapses (Salter and Beggs, 2014; Schafer et al., 2012).

However, the relative contribution of neuronal and astrocytic MAGL to 2-AG signal termination may differ at individual synapses as not all CB1 receptor nerve terminals express MAGL (Monory *et al*., 2006; Uchigashima *et al*., 2011). Thus, the extent of 2-AG-mediated inter-synaptic crosstalk may vary from synapse to synapse depending on the constellation of 2-AG-mediated signaling molecules around synapses and the anatomical organization of presynaptic, postsynaptic, and glial elements.

Binding levels of [^3^H]T-401 in brain sections from a mouse model of temporal lobe epilepsy revealed significant reductions in binding in the hippocampus and cerebral cortex and in subcortical areas. The reduction in [^3^H]T-401 binding observed mostly in the ipsilateral hippocampus suggests a specific effect of the KA treatment on MAGL in the epileptic focus. Additionally, the analysis of cortical subparts showed a decline in [^3^H]T-401 binding in both frontal and temporal cortex, indicating a widespread impact on MAGL activity in these brain regions in the chronic epileptic phase in this model. Whether such changes occur in human epilepsy is still speculative.

Seizures lead to increased hippocampal levels of 2-AG and anandamide (Farrell *et al*., 2021; Marsicano et al., 2003; Wallace et al., 2003; Wettschureck et al., 2006). This may be a result of MAGL reduction occurring within the first days after status epilepticus (Mikkelsen *et al*., 2024). As a consequence, 2-AG activating CB1 expressed in hippocampal excitatory neurons is protective (Katona and Freund, 2008; Kawamura et al., 2006; Monory *et al*., 2006).

Along the same lines, the MAGL inhibitor CPD-4645 reduced spike frequencies and shortening status epilepticus in mice intracerebrally injected with kainic acid (Terrone *et al*., 2018), and another inhibitor ABX-1431 had anticonvulsant effect against hyperthermia-induced seizure in an animal model of Dravets syndrome (Anderson et al., 2022). Finally, JZL184 and JJKK-048 delayed the development of generalized seizures, decreased seizures, and reduced after-discharge duration in the kindling model of temporal lobe epilepsy, but caused only modest effects in fully kindled mice via the CB1R (Griebel *et al*., 2015; von Ruden *et al*., 2015; Zareie *et al*., 2018).

The lower MAGL in chronic epilepsy supports that 2-AG transmission is enhanced not only in the epileptic focus, but also more widespread in the cerebral cortex and beyond. This mechanism represents an endogenous mechanism that reduce excitation in the epileptic brain, because CB1 antagonists increase both seizure duration and frequency (Wallace *et al*., 2003).

Seizures induced by kindling influenced enzyme activity in the amygdala (Colangeli et al., 2023). Masciano et al (2003) showed that mice lacking CB1 in principal forebrain neurons but not interneurons had more excessive seizures after KA (Marsicano *et al*., 2003).

2-AG is a retrograde messenger that regulate synaptic transmission and plasticity (Alger, 2002; Chen et al., 2011; Gao et al., 2010; Kano et al., 2009; Makara et al., 2005; Pacher et al., 2006; Pan et al., 2009; Stella et al., 1997). Thus, it seem likely that 2-AG synthesized in neurons is responsible for synaptic activity while 2-AG synthesized in astrocytes is responsible for modulation of inflammation presumably via CB2 (Mecha et al., 2015). It is likely that the effect of MAGL inhibition might be more determined by the microenvironment rather than the cell type. This intracellular communication is further emphasized by the fact that MAGL inhibition in astrocytes affects microglia gene expression and function (Zhu *et al*., 2023). Anti-convulsive effects of MAGL inhibitors can be blocked by CB1 antagonist suggesting a neuronal effect (Sugaya *et al*., 2016; von Ruden *et al*., 2015; Zareie *et al*., 2018). Nevertheless, MAGL inhibitors had anti-convulsive effects in CB1 KO mice, indicating that also non-CB1 receptors presumably in non-neuronal tissue play a role in the anticonvulsive effect (Terrone *et al*., 2018). To this end, 2-AG antagonizing seizure generation is mediated through CB2, because the CB2 antagonist AM630 produce pro-convulsive effects when co-administrated with the CB receptor agonist WIN55,212-2 (Rizzo et al., 2014; Sugaya *et al*., 2016).

However, the significant increase in [^3^H]PBR28 binding, that reflect TSPO and considered an inflammation marker in the ipsilateral hippocampus of the KA-treated group indicates the presence of neuroinflammation in the epileptic focus. Interestingly, a negative correlation was found between [^3^H]PBR28 binding and [^3^H]T-401 binding, suggesting a potential interaction between neuroinflammation and MAGL activity in the context of epilepsy.

In conclusion, the present study provides insights into the binding characteristics of the MAGL radiotracer [^3^H]T-401 in human and mouse brain tissue. The results support the use of [^3^H]T-401 as a reliable tool for studying MAGL binding in both preclinical and clinical settings. The observed alterations in [^3^H]T-401 binding in the mouse model of temporal lobe epilepsy suggest a potential involvement of MAGL in the pathophysiology of epilepsy. The correlation between [^3^H]PBR28 binding and [^3^H]T-401 binding indicates a potential interplay between neuroinflammation and MAGL activity in epilepsy. These findings pave the way for further investigations into the therapeutic implications of modulating MAGL activity in CNS disorders.

## Author contributions

SSA, BAP, JFB, and JDM planned the experiments. KBL and LP recruited the patients, provided clinical input and conducted neuropathological examination of the resected tissues. BBA provided radiotracers and other molecules. FG and JFB planned the animal experiments and organized collection of the mouse brains. SSA, BAP and JDM conducted the autoradiography experiments, analysed the data by imaging software, and produced the figures. JDM wrote the first draft of the paper, and all authors provided feedback and approved the final version before submission.

## Acknowledgements.

The authors thank Venceslas Duvaue at Synapcell, France for diligent generation of the intra-hippocampal KA mouse model tissue and supplying mouse seizure frequency data. The work was supported by the NOVO Nordisk Foundation (grant no NNF23OC0081536).

## Ethical

The study was approved by the Ethical Committee in the Capital Region of Denmark (H-2-2011-104) and written informed consent was obtained from all patients before surgery. All in-life in-vivo experiments were conducted at SynapCell’s facility. They were approved by the ethical committee of the Grenoble Institute of Neuroscience, University Grenoble Alpes, and performed in accordance with the European Committee Council directive of September 22, 2010 (2010/63/EU) under the responsibility of Dr. Céline Ruggiero (Licence #: 38 11 49).

